# Histidine betaine trimethylammonia-lyase, a novel enzyme coupled with terminal urocanate reductase in *Shewanella woodyi* grown anaerobically

**DOI:** 10.1101/2025.08.13.670108

**Authors:** Yulia V. Bertsova, Marina V. Serebryakova, Alexander A. Baykov, Alexander V. Bogachev

## Abstract

Bacteria coping with oxygen deficiency can switch to alternative terminal electron acceptors, which can be normal metabolic intermediates or products of dedicated coupled reactions. In the latter case, the genes for the respective terminal reductase and coupling enzyme are expected to cluster in the genome. Here, we determined the roles of two uncharacterized periplasmic proteins encoded by the *swoo_3912*–*swoo_3913* gene cluster in the facultatively anaerobic marine bacterium *Shewanella woodyi*. We confirmed the current database annotation of the former protein as “urocanate reductase” but identified the latter protein as a previously unknown histidine betaine trimethylammonia-lyase (HBTL) enzyme. HBTL converts histidine betaine into urocanate and trimethylamine and is remarkably specific for histidine betaine as substrate. HBTL requires Mg^2+^ for activity and undergoes slow reversible inactivation at low Mg^2+^ concentrations. HBTL activity was not evident in *S. woodyi* cells grown aerobically but was induced in cells grown anaerobically. Both histidine betaine and urocanate supported anaerobic *S. woodyi* growth and, hence, respiration. Similar gene clusters are found in many anaerobic bacteria, suggesting a wide occurrence of the anaerobic respiration pathway discovered in this work in the bacterial world.

## 1. Introduction

Anaerobic heterotrophs can cope with low energy yield of fermentation by recruiting alternative terminal electron acceptors for anaerobic respiration—inorganic (sulfate, nitrate, Fe^3+^, and others) [1] or organic compounds (fumarate, acrylate, urocanate, itaconate, and various cinnamate derivatives) [2–6]. Relatively high redox potentials of organic compounds make them less favorable in comparison with the inorganic electron acceptors, O_2_ in particular. However, inorganic acceptors are not always easily available, whereas the organic acceptors are either the products of the central metabolism or can be produced by simple transformations of common secondary metabolites. Thus, fumarate can be formed from malate, aspartate, or tartrate [7], whereas acrylate is a product of the osmolyte dimethylsulfonylpropionate transformation [5,8].

The diversity of natural organic terminal electron acceptors, as well as the pathways for their synthesis by anaerobic organisms, have not yet been fully determined. In bacteria, anaerobic respiration based on the organic acceptors is often carried out by so-called flavocytochromes [4,9]. These proteins are found on the outer surface of the cytoplasmic membrane and consist of two domains. The FAD-binding domain (*FAD_binding_2*, Pfam ID: PF00890) binds and reduces the organic electron acceptor. The heme C-containing domain (a multiheme cytochrome *c*) or, alternatively, a domain containing covalently-bound FMN (*FMN_bind*, PF04205) carries electrons to the FAD-containing domain from an external source of reducing equivalents. The two domains may exist within a single polypeptide or form two distinct subunits.

In the genomes of many bacteria, flavocytochrome gene(s) forms a cluster with the gene encoding another putative periplasmic enzyme, often annotated as “aromatic amino acid ammonia-lyase” (Fig. S1). The latter enzyme catalyzes the removal of an amino group from histidine, phenylalanine, or tyrosine to yield urocanic, cinnamic, or *p*-coumaric acids, respectively [10]. This homotetrameric enzyme contains a unique electrophilic prosthetic group, MIO, generated autocatalytically from a conserved Ala-Ser-Gly sequence [11]. The group is directly involved in α-amino group withdrawal from substrates and is essential for catalysis [12]. Synchronously, a proton is removed from the β-carbon by conserved Tyr residue, resulting in the formation of a *trans*-isomer of the corresponding α,β-enoic acid [10]. Considering that the α,β-enoic acids can function as terminal electron acceptors [3], we hypothesized that the above gene cluster encodes for components of a yet unknown metabolic pathway of terminal electron acceptor formation and its use for anaerobic respiration.

In this study, we tested this hypothesis with a facultative anaerobic marine bacterium *Shewanella woodyi* [13]. Its genome contains adjacent *swoo_3912* and *swoo_3913* genes (UniProt IDs: B1KFJ5and B1KFJ6), whose protein products are annotated solely based on their sequences, as “urocanate reductase” and “aromatic amino acid ammonia-lyase” in the UniProt and Kegg databases. We have isolated these proteins for the first time and determined their true substrate specificities, kinetic characteristics, and roles in anaerobic respiration. The results support our hypothesis but also indicate that one of the studied genes encodes for a novel enzyme, histidine betaine trimethylammonia-lyase, involved in anaerobic respiration.

## 2. Materials and Methods

### 2.1 Bacterial strains and growth conditions

*S. woodyi* DSM 12036 cells were grown aerobically or anaerobically at 25°C in an MR medium containing 3.15% artificial sea salt, 20 mM L-lactate, 0.5% peptone, 0.25% yeast extract, and 20 mM HEPES-NaOH (pH 7.5). Where required, the growth medium was supplemented with 10 mM histidine betaine (HB), urocanate, histidine, trimethylamine hydrochloride, or dimethyl sulfoxide (DMSO). Anaerobic cultivations were carried out in sealed glass flasks that were completely filled with the medium.

### 2.2 Genetic manipulations

To construct a plasmid encoding a 6×His-tagged variant of Swoo_3912, the *swoo_3912* gene was amplified from the genomic DNA of *S. woodyi* by PCR using high-fidelity Tersus polymerase (Evrogen) and 5’-<u>CCATGG</u>GTTTCAAATTTAAAAAGTCAG / 5’-<u>GAATTC</u>TTGCTTAGCCGCTTCTTG primer pair (restriction sites for *Nco*I and *Eco*RI are underlined). The amplified 1766 bp fragment was cloned into a pBAD-TOPO vector (Invitrogen) to generate the pTB_URD plasmid. The direct subcloning of *swoo_3912* into pBAD/*Myc*-His C vector (Invitrogen) using the introduced *Nco*I and *Eco*RI sites was hampered by the intrinsic *Nco*I site in the *swoo_3912* sequence. Therefore, we subcloned the 715 bp *Nco*I-*Hinc*II fragment from pTB_URD plasmid into the pBAD/*Myc*-His C vector treated by *Nco*I and *Pvu*II endonucleases (resulting in pBAD_715 plasmid). Further, we subcloned the 1076-bp fragment from pTB_URD plasmid into the pBAD_715 plasmid using the *Sac*II/*Eco*RI sites, resulting in the plasmid pBAD_URD, encoding full-length Swoo_3912 with a C-terminal 6×His-tag. The Swoo_3912-encoding region of pBAD_URD was verified using DNA sequencing, and the plasmid was electroporated into *Vibrio cholerae* O395N1 *tox*T::*lac*Z cells.

To construct the plasmid encoding cytoplasmic 6×His-tagged variant of Swoo_3913, the 69–1659 nucleotides of the *swoo_3913*gene missing those for N-terminal leader peptide were amplified from the genomic DNA of *S. woodyi* by PCR using Tersus polymerase and the forward/reverse primers 5’-<u>CATATG</u>GATACTATCAGCCTAACC /5’-<u>CTCGAG</u>GTATTTCATTAAGAATTT (restriction sites for *Nde*I and *Xho*I are underlined). The amplified 1596-bp fragment was cloned into a pCR4-TOPO vector (Invitrogen), resulting in a pCR_HAL plasmid. Further, we subcloned *swoo_3913*into the pSCodon vector (Delphi Genetics) using the *Nde*I/*Xho*I sites, resulting in the plasmid pSC_HAL. DNA sequencing was used to verify the Swoo_3913-encoding region of pSC_HAL, and the plasmid was transformed into the *Escherichia coli* BL21 (DE3) strain.

### 2.3 Swoo_3912 and Swoo_3913 isolation

Recombinant 6×His-tagged Swoo_3912 was heterologously produced in the periplasm of *V. cholerae* cells and purified from the membrane fraction of this bacterium using metal chelate chromatography, as described previously [4]. Swoo_3912 concentration was determined using the extinction coefficient ε_452-600_ of 26.3 mM^-1^ cm^-1^ [4].

6×His-Tagged Swoo_3913 was produced in the cytoplasm of *E. coli /* pSC_HAL cells. The cells were grown to the late-exponential phase (*A*_600_ ≈ 1.5) in LB medium at 37°C, after which Swoo_3913 synthesis was induced with 0.2% (w/v) lactose, and the cells were further grown for 16 h at 15°C. Swoo_3913 was purified from *E. coli* cells using metal chelate chromatography [14]. The protein concentration was determined by the bicinchoninic acid method [15] using bovine serum albumin as the standard. Stock solution of Swoo_3913 was stored frozen in 20 mM Tris-HCl buffer (pH 8.0) with 20 % glycerol added and was diluted 100-fold with the same buffer before use. Purified Swoo_3912 was stored frozen in the same medium with 0.05% *n*-dodecyl β-D-maltoside added.

### 2.4 Mass-spectral analysis

The procedures for MALDI-TOF MS analysis of the purified proteins have been previously described [16]. Proteins were separated by SDS-PAGE and in-gel digested with trypsin. The resulting peptides were analyzed using an UltrafleXtreme MALDI-TOF-TOF mass spectrometer (Bruker Daltonik).

### 2.5 Swoo_3912 activity assay

The reductase activity of Swoo_3912 was determined by following the oxidation of reduced methyl viologen (MV, ε_606_ = 13.7 mM^-1^ cm^-1^) using a Hitachi-557 spectrophotometer. The assay was performed in a 3.2 mL airproof cuvette at 25°C. The standard assay mixture contained 100 mM Tris-HCl (pH 8.0), 1 mM MV, and 0.1 mM electron acceptor. Anaerobic conditions and MV reduction were achieved by gradually adding dithionite until the absorbance at 606 nm reached ∼1.5, which corresponds to ∼100 μM reduced MV and ∼900 μM oxidized MV concentrations in the medium. One unit of enzyme activity was defined as the enzyme amount catalyzing oxidation of 2 μmol MV per 1 min.

The Michaelis-Menten parameters of urocanate reductase reactions were estimated from the kinetics of 0.02–15 μM urocanate reduction, as monitored by absorbance at 606 nm. All measurements were carried out at saturating concentrations of reduced MV, based on the *K*_m_ ≤ 5 μM for MV. Rates were estimated at 30 time points along the complete progress curve as the slopes of the tangents (−*d*[MV]/*d*t) using MATLAB (The MathWorks, Inc., USA). The residual substrate (electron acceptor) concentration was calculated at each point from *A*_606_ using the MV:acceptor stoichiometry of 2:1 and assuming that the limiting value of *A*_606_ corresponds to 100% conversion of the electron acceptor. This procedure yielded more reliable rate values at (sub)micromolar substrate concentrations than the traditional initial-velocity method. The similarity of the *K*_m_ values thus determined at different concentrations of Swoo_3912 (0.06-0.24 μg/mL) and urocanate (5-15 μM) indicated no interference from possible product inhibition, reverse reaction contribution, and enzyme activation/inactivation in the course of the reaction, the common obstacles in integral kinetics analysis [17,18]. The Michaelis-Menten equation was fitted to the rate data using nonlinear regression analysis.

### 2.6 Swoo_3913 activity assay

L-Histidine, L-phenylalanine, L-tyrosine, and L-tryptophan ammonia-lyase activities of Swoo_3913 were tested with an Aminco DW-2000 spectrophotometer by following the formation of urocanate (277 nm), cinnamate (269 nm), *p*-coumarate (286 nm), or 3-indoleacrylate (340 nm), respectively. The assay mixture contained medium A (20 mM Tris-HCl, pH 8.5, and 1.75 % marine salts), Swoo_3913 (20 μg/mL), and an L-amino acid (1 mM histidine, 1 mM phenylalanine, 0.1 mM tyrosine, or 0.1 mM tryptophan). The same assay was used to test the lyase activities of Swoo_3913 with D-histidine, L-imidazolelactate, and L-carnosine by following the formation of urocanate at 277 nm.

The ammonia-lyase activities of Swoo_3913 with all other proteinogenic amino acids (except cysteine) were measured using Nessler’s reagent. The assay mixture (200 μL) contained 2 mM amino acid and purified Swoo_3913 (50 μg/mL) in medium A. The reaction mixture was incubated at 30°C for 4 h, and the reaction was stopped by adding trifluoroacetic acid to 5% concentration. The mixture was centrifuged at 11,000 *g* for 2 min to sediment the precipitate.

The supernatant was neutralized by adding 800 μL of 100 mM Na_2_CO_3_ containing 50 mM Na,K-tartrate and mixed with 1 mL of Nessler’s reagent. The sample was incubated at room temperature for 10 min, and its absorbance was measured at 425 nm. The trifluoroacetic acid was added before the enzyme in the control samples.

Histidine betaine (HB) and ergothioneine trimethylammonia-lyase activities of Swoo_3913 were measured with an Aminco DW-2000 spectrophotometer by following the formation of urocanate (at 277 nm, ε = 18.8 mM^-1^ cm^-1^) or thiourocanate (at 311 nm, ε = 22.5 mM^-1^ cm^-1^), respectively. Activity values refer to the initial substrate conversion velocities. The assay mixture contained medium B (50 mM Bis-Tris propane-HCl, pH 8.5, and 10 mM MgSO_4_), Swoo_3913 (0.1–10 μg/mL), and 0.01–2 mM substrate. The pH-dependence of the HB trimethylammonia-lyase activity of Swoo_3913 was measured in a buffer containing 100 mM Bis-Tris propane-HCl, 10 mM MgSO_4_, and 1 mM HB. The rate values (*v*) were analyzed using the following equation:

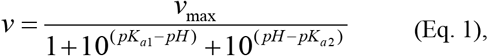

where *v*_max_ is the limiting value of *v* and *pK*_a1_ and *pK*_a2_ are the acid dissociation constants for two catalytically important groups.

### 2.7 Determination of the activity of histidine betaine trimethylammonia-lyase in S. woodyi cells

*S. woodyi* cells were grown aerobically or anaerobically with an appropriate potential electron acceptor, harvested by centrifugation (11,000*g*, 2 min), and washed twice with the medium containing 10 mM Tris-HCl (pH 8.0), 0.5 M NaCl, and 5 mM MgSO_4_. HB trimethylammonia-lyase activity in cells was determined spectrophotometrically at 277 *versus* 312 nm (ε = 17 mM^-1^ cm^-1^) in the medium B containing 1 mM HB.

## 3. Results

### 3.1 Swoo_3912 and Swoo_3913 isolation

The genes for 6×His-tagged full-size Swoo_3912 (including its leader peptide) and truncated Swoo_3913 (lacking its leader peptide) were expressed in *V. cholerae* and *E. coli*, respectively. The choice of *V. cholerae* cells was explained by their ability to flavinylate foreign periplasmic proteins [4]. The recombinant Swoo_3912 and Swoo_3913 proteins were isolated by metal affinity chromatography with the yields of 0.1 and 70 mg per 1 L of bacterial culture, respectively.

The final preparation of Swoo_3912 demonstrated several protein bands on SDS-PAGE analysis. One of them strongly fluoresced when illuminated at 473 nm, indicating a covalently bound flavin [19]. MALDI-MS analysis of this band confirmed its identity with Swoo_3912 (sequence coverage: 72%; Fig. S2). The major non-fluorescent bands were identified as Swoo_3912 fragments because their peptide spectrum did not contain any peptides generated from the N-terminal domain of the protein. A likely corollary is that the heterologously produced Swoo_3912 is only partially flavinylated and is prone to degradation in the non-flavinylated state.

In contrast, Swoo_3913 was isolated as an electrophoretically pure ∼51 kDa full-size protein (Fig. 1). MALDI-MS analysis confirmed its protein identity (sequence coverage: 80%; Fig. S3).

**Fig. 1.**
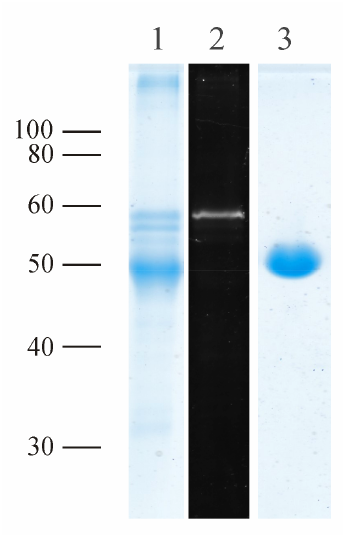
SDS-PAGE of isolated Swoo_3912 (lanes *1* and *2*) and Swoo_3913 (lane *3*). The gels were stained with Coomassie Blue (lanes *1* and *3*) or scanned with a Typhoon FLA 9500 laser scanner (GE Healthcare) with excitation at 473 nm and the detection of emission using the SYBR Green II protocol (lane *2*). The protein load was 4 μg per lane. The bars with numbers on the left side denote the positions and molecular masses of marker proteins.

### 3.2 Structure-based prediction of the enzymatic identities of Swoo_3912 and Swoo_3913

The *S. woodyi* protein Swoo_3912 is a close homolog of *S. oneidensis* urocanate reductase *So*_UrdA [4] (70% identity, 86% similarity). Both proteins are formed by *FMN_bind* and *FAD_binding_2* domains and contain additionally an N-terminal Sec-type lipoprotein signal peptide. Furthermore, both proteins contain a flavinylation motif (DGITGA**T** in Swoo_3912 and DGIAGA**T** in *So*_UrdA; the residue to which FMN is covalently linked shown in bold [19,20]) and conserved residues involved in urocanate carboxyl and imidazole nitrogen binding [21,22] (Fig. S4). Thus, the Swoo_3912 protein is highly likely an urocanate reductase, in accordance with its annotation in the UniProt and Kegg databases.

Considering that urocanate is a product of histidine deamination, the genetically coupled Swoo_3913 protein could be a histidine ammonia-lyase that catalyzes the deamination reaction. The amino acid sequence of this protein demonstrates moderate similarity with that of the characterized histidine ammonia-lyase of *Pseudomonas putida* (26% identity, 42% similarity; Fig. S5). The Swoo_3913 protein contains most conserved catalytic residues of histidine ammonia-lyases [10], including Gln308, Arg314, and Asn339, which are involved in histidine carboxylate binding, His117 and Glu467, involved in imidazole N_τ_ and N_π_ binding, and the substrate β-proton acceptor Tyr77 (Fig. S5). However, Swoo_3913 lacks the Ala-Ser-Gly sequence that participates in the autocatalytic formation of the prosthetic electrophilic group MIO in aromatic amino acid lyases and mutases [10,11]. MIO interacts with the α-amino group of the substrate during its withdrawal by ammonia-lyase [12]. At this stage, the sequence analysis suggested that the Swoo_3913 substrate, if any, may be a histidine analog with the α-amino group replaced by a different function.

To further narrow our search among numerous naturally occurring histidine analogs, we compared the 3D structure of Swoo_3913 (Fig. S6 A,B) predicted by AlphaFold [23] with the known protein structures available in the PDB. The DALI server [24] was used for the comparison as the DALI algorithm allows detection of similarity even with phylogenetically distant homologues. The best match (z-score = 43, r.m.s.d. = 2.3 Å) was found with *Treponema denticola* ergothionase (*Td*_ETL, PDB Id: 6s7j) [25] (Fig. S6C). Ergothionases are involved in the catabolism of ergothioneine, a natural antioxidant wide-spread in the biosphere [26] (Fig. 2), and catalyze trimethylamine removal from ergothioneine to form thiourocanate (Fig. 2) [27,28]. Importantly, trimethylamine is a much better leaving group than the amino group, explaining why ergothionases, unlike histidine ammonia-lyases, do not need a MIO prosthetic group for catalysis [25,28]. Thus, ergothioneine or a derivative thereof may be a potential Swoo_3913 substrate.

**Fig. 2.**
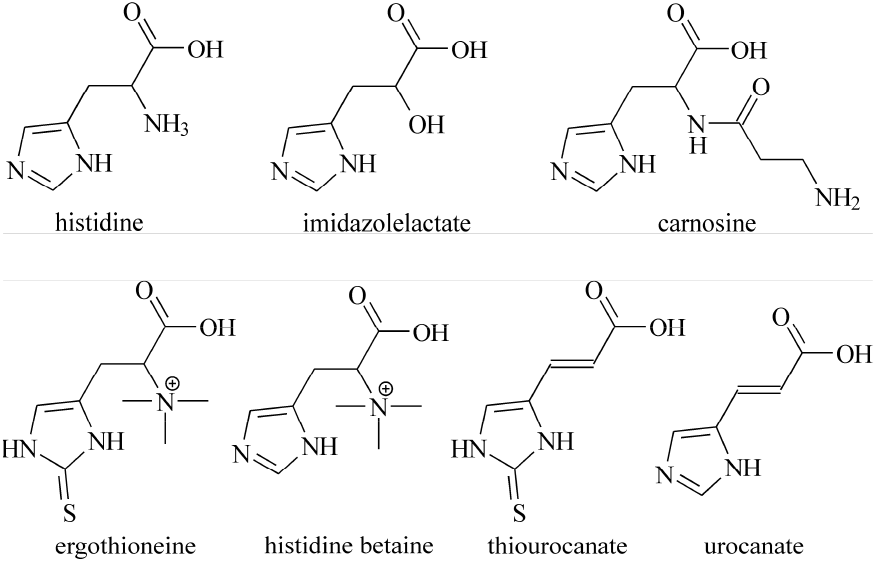
Histidine and its derivatives considered in this study.

This prediction was further refined by amino acid sequence comparison between Swoo_3913 and the ergothionase *Td*_ETL (Fig. S5). The comparison indicated that Swoo_3913 contains nearly all the amino acid residues involved in ergothionase catalysis [25], with the exception of *Td*_ETL Lys384 being replaced by a neutral Asn439 residue in Swoo_3913.

Lys384 allows ergothioneine conversion into its thiolate form and thiolate group binding and is essential for Td_ETL ergothionase activity [25]. Based on the above, one could expect the Swoo_3913 substrate be an ergothioneine derivative with no sulfur atom in its structure, i.e. histidine betaine (hercynine) (HB, Fig. 2).

### 3.3 Experimental verification of the structure-based predictions

With methyl viologen as the electron donor, the isolated Swoo_3912 catalyzed urocanate reduction (Fig. 3) with *k*_cat_ of 65 ± 4 s^-1^ and *K*_m_ of 0.7 ± 0.1 μM. The enzyme exhibited no activity with fumarate, cinnamate, caffeate, methacrylate, and acrylate and only marginal activity (≤ 0.08 s^-1^) with thiourocanate. Thus, Swoo_3912 is a highly selective catalyst, with the specificity consistent with the above sequence-based prediction (section 3.2.) and database annotation as urocanate reductase *Sw*_UrdA.

**Fig. 3.**
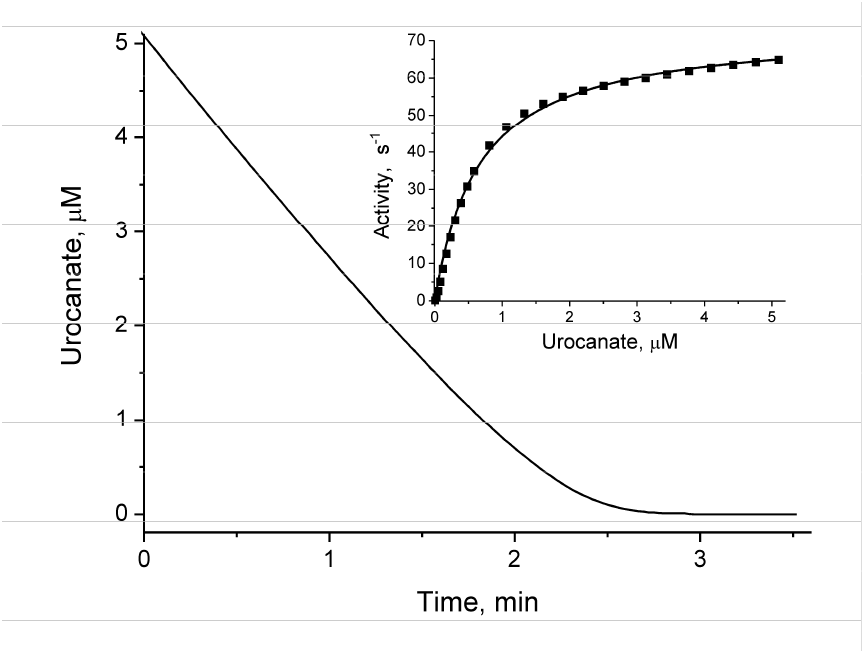
Time course of urocanate reduction by Swoo_3912. Inset: Dependence of the *Swoo*_3912 activity, as estimated from the tangents to the reaction time course, on residual urocanate concentration; the line shows the best fit of the Michaelis-Menten equation.

Identification of the enzymatic activity of Swoo_3913 was not that easy. This protein demonstrated no histidine ammonia-lyase activity (did not deaminate L-or D-histidine) or a same activity against any other proteogenic amino acids (cysteine was not tested because it interferes with the ammonia assay used). Thus, the observed inability of Swoo_3913 to deaminate aromatic amino acids (histidine, phenylalanine, tyrosine, and tryptophan) is inconsistent with its annotation as “phenylalanine/histidine ammonia-lyase” in GenBank, UniProt, and Kegg.

No activity was similarly detected with the histidine analogs carnosine and imidazolelactate, indicating that a blockage or mere absence of the substrate amino group is not enough for histidine analog being a sunstrate of Swoo_3913. Ergothioneine was converted at a low but measurable rate (0.05 s^-1^), suggesting that the trimethylamino group had a stimulatory effect.

Finally, and fully consistent with the structure-based predictions, high activity of Swoo_3913 was observed with histidine betaine (Fig. 4A), which differed from ergothioneine by lacking its thio group. Thus, the Swoo_3913 protein represents a novel lyase enzyme, histidine betaine trimethylammonia-lyase (HBTL), that converts histidine betaine to yield *trans*-urocanate and trimethylamine. This is a novel reaction that seemingly occurs in the living world.

**Fig. 4.**
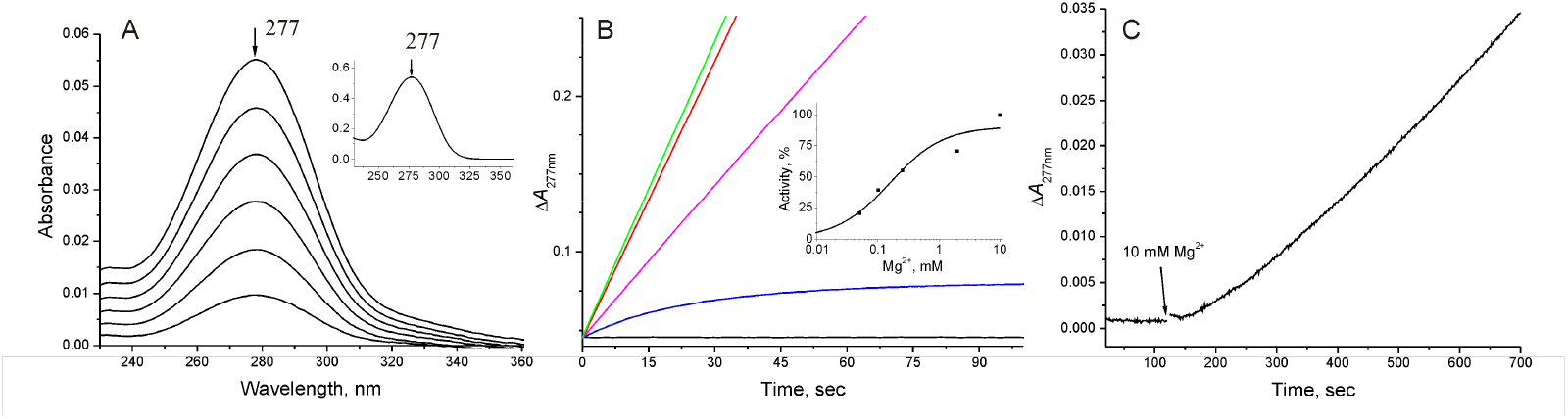
The histidine betaine trimethylammonia-lyase activity of Swoo_3913. (A) Spectral changes in the UV region during incubation of Swoo_3913 (0.08 μg/mL) with 0.5 mM HB. The spectra were recorded in 30-s intervals, starting 30 s after reaction onset. The inset shows the spectrum of 30 μM *trans*-urocanate. The arrows indicate the maximum wavelength (277 nm). The assay medium contained 50 mM Bis-Tris propane-HCl (pH 8.5) and 10 mM MgSO_4_. (B) The effect of Mg^2+^ ions on the HBTL activity of Swoo_3913 (1.1 μg/mL) in the medium containing 50 mM Bis-Tris propane (pH 8.5) and 0.5 mM HB. Further additions: green curve, 1.75% marine salt; red curve, 10 mM MgSO_4_; magenta curve, 2 mM MgSO_4_; blue curve, 50 μM MgSO_4_; black curve, no additions. The reaction was started with the enzyme. Inset: Dependence of the initial HBTL activity on Mg^2+^ concentration. The curve shows the best-fit binding isotherm with *K*_0.5_ of 0.16 ± 0.04 mM. (C) Reactivation of Swoo_3913 (0.08 μg/mL) preincubated for 120 s with 50 mM Bis-Tris propane-HCl (pH 8.5) and 0.5 mM HB before adding 10 mM MgSO_4_ (indicated by the arrow).

### 3.4 Kinetic characterization of Swoo_3913 histidine betaine trimethylammonia-lyase activity

Initial measurements of Swoo_3913 activities were performed in the pH 8.5 buffer containing artificial sea salt to mimic the *in vivo* periplasmic composition of the marine bacterium *S. woodyi* (Fig. 4B). No HBTL activity was observed in the absence of the sea salt (Fig. 4B). However, further experiments indicated that Mg^2+^ ions alone at a concentration of 10 mM could fully replace the sea salt as the essential activator (Fig. 4B). Ca^2+^ ions could also activate Swoo_3913 to the same level but at approximately six fold higher concentrations in comparison with Mg^2+^ ions.

The linear reaction time courses obtained in the presence of 10 mM Mg^2+^ and excess of substrate (Fig. 4B) indicated that the enzyme was stable in these assay conditions. At a low Mg^2+^ concentration (0.05 mM), enzyme activity was much lower and decreased in time, resulting in a concave time course. This apparent enzyme inactivation was slowly but fully reversed by incubation with 10 mM Mg^2+^ (Fig. 4C). Interestingly, no inactivation was observed when Swoo_3913 was incubated for hours at low or zero Mg^2+^ concentration in the absence of the substrate histidine betaine. These findings suggest that substrate binding to Swoo_3913 in the absence of Mg^2+^ results in a tight dead-end complex, whose further conversion requires metal ion cofactor binding, which occurs slowly because the substrate shields the cofactor-binding site from medium Mg^2+^. In the normal catalytic cycle, the metal cofactor apparently binds before the substrate. Alternatively, Mg^2+^ may be required for product release from the active site or may convert the enzyme into the active conformation or oligomeric state. No such effect of metal ions has been previously documented for aromatic amino acid ammonia-lyases or ergothionases. This unique property of Swoo_3913 may be associated with the host bacterium’s alkali-earth metal cation-reach marine habitat.

The substrate concentration dependence of the Swoo_3913 HBTL activity was hyperbolic (Fig. 5A), yielding the following Michaelis-Menten parameters: *K*_m_ = 103 ± 8 μM and *k*_cat_ = 18 ± 1 s^-1^, which are comparable with the values reported for *Td*_ETL (*K*_m_ = 46 μM and *k*_cat_ = 64 s^-1^ [25]). Similar parameter values were obtained for Swoo_3913 using histidine betaine prepared by L-histidine methylation with iodomethane [29] in this lab or obtained from Shanghai Macklin Biochemical Technology Co., Ltd. (Shanghai, China).

**Fig. 5.**
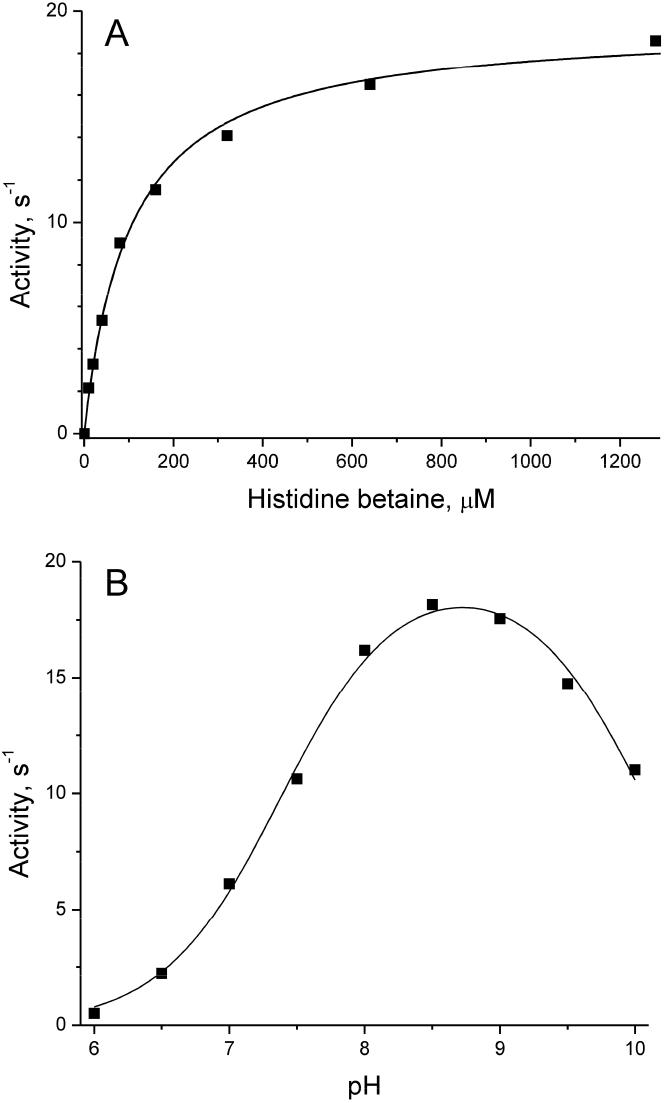
Dependence of Swoo_3913 activity on HB concentration at pH 8.5 (A) and on pH at 1 mM HB concentration (B). The lines show the best fits of the Michaelis-Menten equation and Eq. 1, respectively.

The pH dependence of the Swoo_3913 activity was bell-shaped and seemingly symmetrical, with a maximum at alkaline pH (8.6) and the *pK*_a_ values for two catalytically important ionizable groups of 7.4 and 10.1. These groups likely refer to the catalytic step (*k*_cat_), as the substrate concentration used was nearly saturating, at least at pH 8.5 (Fig. 5A). The observed *pK*_a_ values are approximately one unit greater than those reported for *Td*_ETL [25].

### 3.5 Physiological role of Swoo_3912 and Swoo_3913 in S. woodyi

Oxygen deficiency favors the induction of enzymatic activities that use terminal electron acceptors other than oxygen, provided that the appropriate substrate is present in the medium [4– 6,30]. To define the role of Swoo_3913 in anaerobic respiration, we determined the effects of growth conditions on the HBTL activity in *S. woodyi*. The bacterium grown aerobically in the absence or presence of HB or urocanate demonstrated no such activity, whereas its level was as high as ∼60 nmol/min per 1 mg cell protein in the bacterium grown anaerobically (Table 1).

**Table 1.** Histidine betaine trimethylammonia-lyase activity of *S. woodyi* cells grown under different conditions (in nmol/min per 1 mg of cell protein)

| Growth conditions | Activity<sup>a</sup> |
| --- | --- |
| +O<sub>2</sub> | < 1 |
| +O<sub>2</sub> + 10 mM urocanate | < 1 |
| +O<sub>2</sub> + 10 mM HB | < 1 |
| -O<sub>2</sub> | 59 ± 7 |
| -O<sub>2</sub> + 10 mM DMSO | 64 ± 9 |
| -O<sub>2</sub> + 10 mM urocanate | 67 ± 6 |
| -O<sub>2</sub> + 10 mM HB | 69 ± 7 |
| -O<sub>2</sub> + 10 mM trimethylamine hydrochloride | 53 ± 7 |

<sup>a</sup> Mean value ± SD for three independent replicates.

These findings suggest that Swoo_3913 plays a role in anaerobic respiration.

Interestingly, HB and urocanate did not significantly increase Swoo_3913 activity in anaerobically grown cells (Table 1). Genes involved in anaerobic respiration usually colocalize with the genes of transcription regulators on the chromosome to establish autoregulatory circuits that concordantly regulate neighboring genes [6]. However, no transcription regulator was found in the immediate vicinity of the *swoo_3912*-*swoo_3913* gene cluster (Fig. S1), possibly explaining the inability of the substrates and products of histidine betaine trimethylammonia-lyase to induce Swoo_3913 activity. Notably, *S. oneidensis* fumarate reductase Fcc_3_, a classical anaerobic respiratory enzyme, also follows the same induction pattern—its induction during anaerobiosis does not require fumarate presence in the growth medium [31]. Hence, the lack of a substrate effect on Swoo_3913 synthesis does not rule out its involvement in anaerobic respiration.

To determine whether *S. woodyi* can use urocanate or its precursor (HB) as the terminal electron acceptor for anaerobic respiration, the bacterium was grown in the presence of these compounds using the classical electron acceptor DMSO as the control. This experiment was complicated by the fact that *S. woodyi* cannot grow in the minimal medium under anaerobiosis, whereas it grows even in the absence of added electron acceptor in reach media [5]. The latter property explains why urocanate and HB stimulated growth yield by only ∼25% (Fig. 6).

**Fig. 6.**
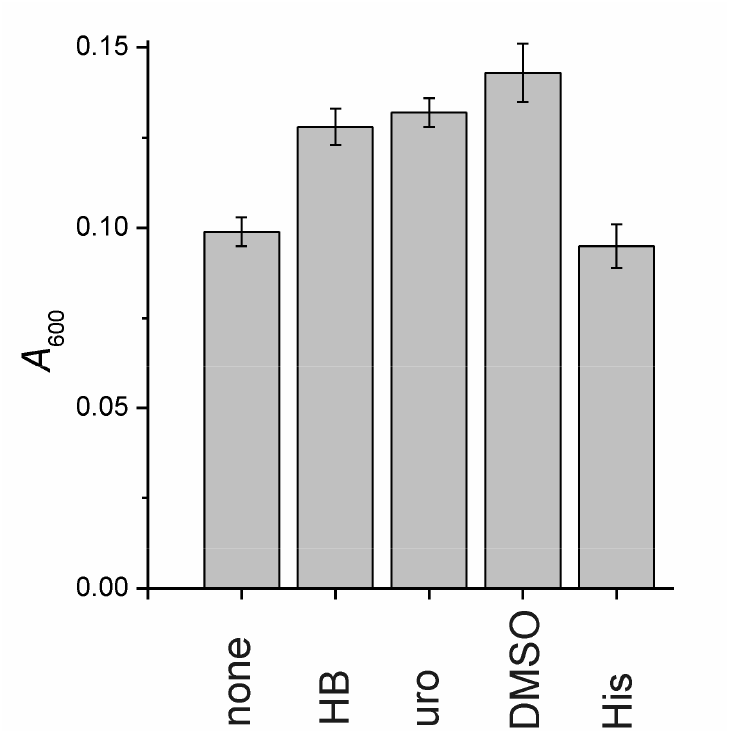
Anaerobic growth of *S. woodyi* in the presence of different electron acceptors. Cells were grown anaerobically for 18 h at 25°C. The growth medium was supplemented with 10 mM HB, urocanate (uro), DMSO, or histidine (His) where indicated. The bars show the final absorbance (light scattering) with the error bars representing SD values in three independent experiments. The initial absorbance of the growth media at 600 nm was 0.005 unit.

However, the effect was comparable with that of DMSO and statistically significant. Thus, both histidine betaine and its urocanate product can support anaerobic respiration in *S. woodyi*. No growth stimulation was caused by histidine, suggesting that the stimulatory effect of HB on *S. woodyi* growth is associated with anaerobic respiration rather than urocanate fermentation.

## 4. Discussion

The above results indicate that the gene cluster *swoo_3912*-*swoo_3913* allows *S. woodyi* to catalyze periplasmic HB conversion into urocanate and use it for anaerobic respiration (Fig. 7). The periplasmic localization of Swoo_3912 and Swoo_3913 indicate that the urocanate generated from HB may be used in the periplasmic space rather than transported into cytoplasm, consistent with the presumed role of urocanate as a terminal electron acceptor for the anaerobic electron transport chain. HB is an ergothioneine precursor and also the product of ergothioneine oxidation when the latter compound functions as an antioxidant [32,33]. That is why high (millimolar) concentrations of HB are generally found in the cytoplasm of ergothioneine-producing organisms (e.g., fungi and cyanobacteria) [34,35]. HB may also play an independent role as an osmolyte (like other betaines [36]) or intracellular pH buffer (like carnosine and its derivatives [37]) in living cells. This explains the widespread occurrence of HB in the environment [38]. Moreover, *S. woodyi* genome contains the gene cluster *swoo_2444*-*swoo_2445*, which encodes the key enzymes of the ergothioneine synthesis pathway, histidine methyltransferase EgtD and sulfoxide synthase EgtB [32]. Thus, it is likely that *S. woodyi* is capable of ergothioneine synthesis and the products of its *swoo_3912*-*swoo_3913* cluster support respiration using own HB that becomes an unnecessary intermediate under anaerobic conditions.

**Fig. 7.**
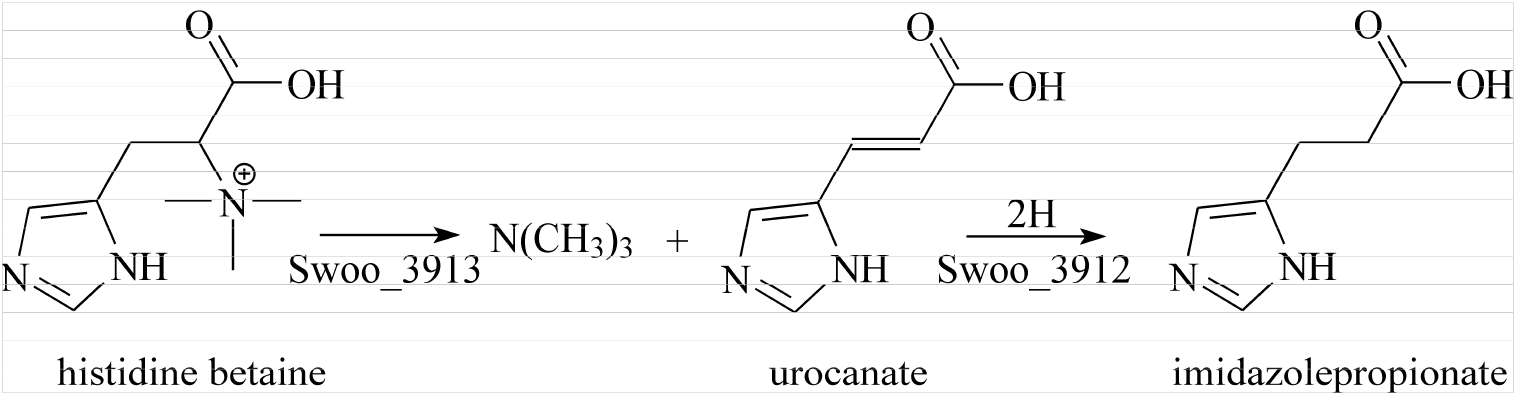
Chemical reactions catalyzed by Swoo_3913 and Swoo_3912 in *S. woodyi* periplasm, resulting in imidazolepropionate (dihydrourocanate) and trimethylamine production from HB via urocanate as an intermediate.

The gene cluster encoding a Swoo_3913-like lyase and flavocytochrome *c* is found in many anaerobic bacteria (Fig. S1), including those colonizing human gut, which suggests a wide occurrence of the anaerobic respiration pathway. However, further experiments are needed to verify the substrate specificity of the corresponding encoded enzymes in different bacteria.

Interestingly, the production of trimethylamine by the gut microbiome is associated with a number of cardiovascular pathologies [39], whereas imidazolepropionate production leads to the development of type 2 diabetes [40] and is associated with many other pathologies [41]. Both compounds are the final products of the aforementioned anaerobic respiration pathway (Fig. 7). Taken together, these findings point to a possible role of this pathway in human gut in the development of the most serious and widespread deceases. Knowing the genes and proteins involved in the biological mechanisms leading to the formation of the causative agents will hopefully help to develop suitable drugs to combat the above deceases.

## CRediT authorship contribution statement

Yulia Bertsova: Methodology, Investigation, Visualization. Marina Serebryakova: Methodology, Investigation. Alexander Baykov: Data curation, Visualization, Writing. Alexander Bogachev: Conceptualization, Methodology, Investigation, Writing, Funding acquisition.

## Acknowledgements

The authors would like to acknowledge Dr. N.S. Melik-Nubarov for the help with histidine betaine synthesis. MALDI MS and laser scanner facilities became available to us in the framework of the Moscow State University Development Program PNG 5.13.

## Data availability

All data are available from the authors at a reasonable request.

