## Supplementary material for "Histidine betaine trimethylammonia-lyase, a novel enzyme coupled with terminal urocanate reductase in *Shewanella woodyi* grown anaerobically": Fig. S

Supplement

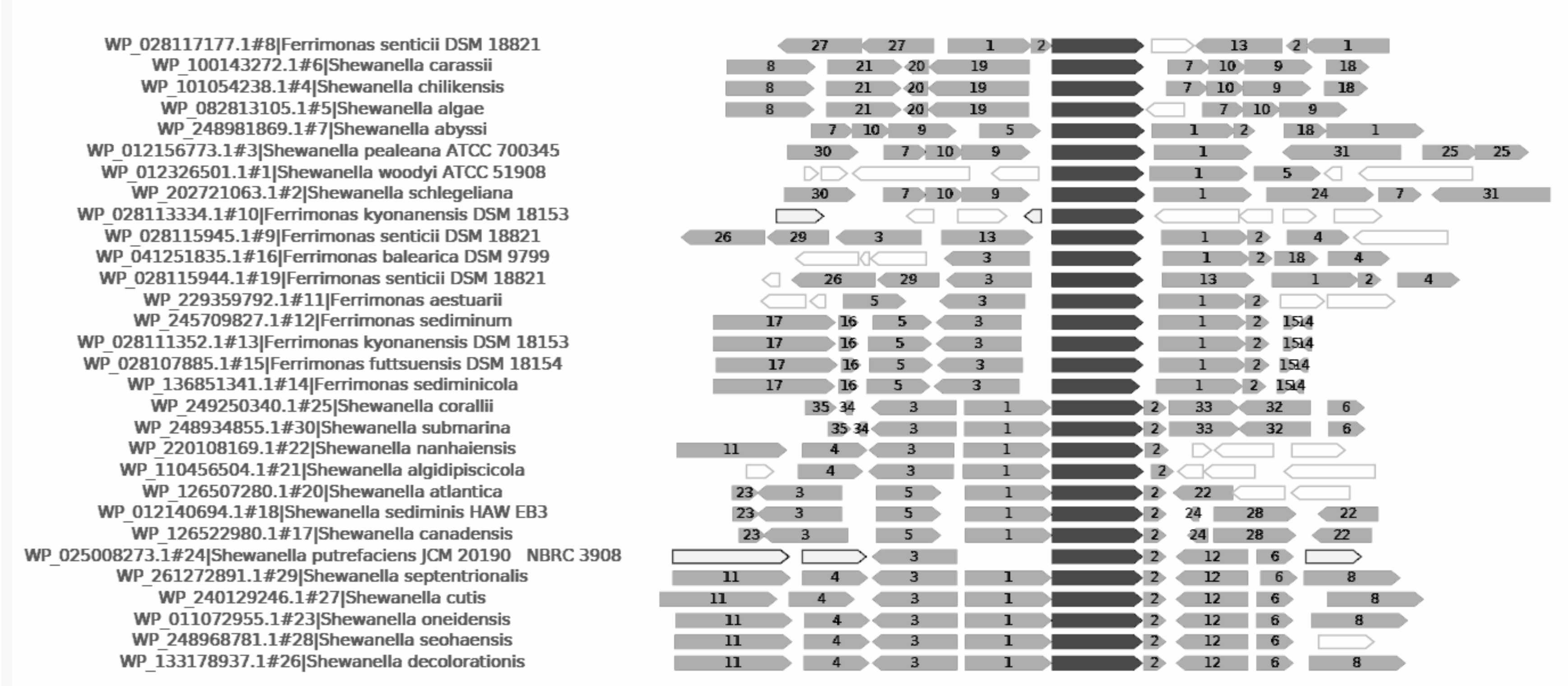


**Fig. S1**. Genomic context analysis of *S. woodyi swoo_3913* gene and its homologs in different *Shewanella* species and related bacteria (black bars) using a webFlaGs service [Saha *et al.*, 2021]. The bacterial species and accession numbers of the corresponding putative ammonia-lyases are shown on the left. The genes encoding homologous proteins are marked by same numbers inside the boxes. The most frequently occurring proteins are as follows: 1, flavocytochrome *c* flavoprotein subunit; 2, multiheme cytochrome *c*; 3, $\sigma_{54}$-dependent Fis family transcriptional regulator; 4, porin; 5, porin domain-containing protein; 6, MBL fold metallo-hydrolase; 7, LuxR family transcriptional regulator; 8, methyl-accepting chemotaxis protein; 9, HAMP domain-containing sensor histidine kinase; 10, response regulator transcription factor; 11, DNA topoisomerase III; 12, $\alpha/\beta$-fold hydrolase; 13, aromatic amino acid ammonia-lyase.

### Mascot Search Results

#### Protein View

Match to: **Swoo_3912_6His** Score: **1420** Expect: **3e-138**
**Swoo_3912_6His**
Found in search of DATA.TXT

Nominal mass ($M_r$): **65472**; Calculated pI value: **5.83**
NCBI BLAST search of Swoo_3912_6His against nr
Unformatted sequence string for pasting into other applications

Variable modifications: Oxidation (M),Propionamide (C)
Cleavage by Trypsin: cuts C-term side of KR unless next residue is P
Sequence Coverage: **72%**

Matched peptides shown in **Bold Red**

```
  1 MGFKFKKSVL GMAVAVPVVL VGCAVTNEMV KSQGTAKGRH GDITVETTFE
 51 NGKITAIDIV KQKENRVLSA AVYKDVKQAI IDNNSINVDG ITGATATSDG
101 LKKAVAKSAE LAGITLVATA AINGKKVNAE PSEYTYDVVV IGAGGAGFSA
151 GVEAVEAGVS AVIIEKQPII GGNSLISGGE MNVAGSWVQQ SMGITDSKAL
201 FIEDTLKGGD YKGDPEMVRV MADNAVAAAE WLRDDIKVDF YKDQIFQFGG
251 HSVKRAVIPK GHTGAEVLSK FAAKAEEIGL PVHMNTTAKN LIQDETGRVV
301 GVKAMKNGKV ITYHAKKAVV MAAGGFGANI EMRKRFNPEY DERYGTTNHA
351 GATGDGIVMS AAINAKTANL GEIQAYPICN PETGAIALIA DARFFGAILV
401 NQEGKRFVEE LDRRDVISNA ILNQTGKYTY VIWNQKIDDL AKTIDMHPGE
451 FNDLHSRGLM FEVESIEEAA AKFNIPLDTL QNTIKDVNQY AATGKDLAFN
501 NRAGLVDMSE GKYWILKATP SVHHTMGGIA TTTKAEVLNN DGDIIKGLYA
551 AGEVTGLTHG SNRLGGNAYT DIIVFGRIAG QEAAKQEFEA YVEQKLISEE
601 DLNSAVDHHH HHH
```

Show predicted peptides also

Sort Peptides By ◉ Residue Number ○ Increasing Mass ○ Decreasing Mass

| Start - End | Observed | Mr(expt) | Mr(calc) | ppm | Miss | Sequence |
|---|---|---|---|---|---|---|
| 40 - 53 | 1547.7494 | 1546.7422 | 1546.7264 | 10 | 0 | R.HGDITVETTFENGK.I (Ions score 101) |
| 78 - 103 | 2616.3537 | 2615.3465 | 2615.3402 | 2 | 1 | K.QAIIDNNSINVDGITGATATSDGLKK.A (Ions score 210) |
| 126 - 166 | 4052.8152 | 4051.8079 | 4052.0630 | -63 | 1 | K.KVNAEPSEYTYDVVVIGAGGAGFSAGVEAVEAGVSAVIIEK.Q (No match) |
| 127 - 166 | 3924.8006 | 3923.7933 | 3923.9680 | -45 | 0 | K.VNAEPSEYTYDVVVIGAGGAGFSAGVEAVEAGVSAVIIEK.Q (Ions score 27) |
| 167 - 198 | 3261.5636 | 3260.5563 | 3260.5806 | -7 | 0 | K.QPIIGGNSLISGGEMNVAGSWVQQSMGITDSK.A (Ions score 185) |
| 167 - 198 | 3277.7136 | 3276.7064 | 3276.5755 | 40 | 0 | K.QPIIGGNSLISGGEMNVAGSWVQQSMGITDSK.A Oxidation (M) (No match) |
| 199 - 207 | 1049.5755 | 1048.5682 | 1048.5804 | -12 | 0 | K.ALFIEDTLK.G (No match) |
| 220 - 233 | 1516.8188 | 1515.8115 | 1515.7504 | 40 | 0 | R.VMADNAVAAAEWLR.D (Ions score 67) |
| 220 - 237 | 1988.0225 | 1987.0152 | 1986.9833 | 16 | 1 | R.VMADNAVAAAEWLRDDIK.V (No match) |
| 238 - 254 | 2015.0366 | 2014.0293 | 2013.9949 | 17 | 1 | K.VDFYKDQIFQFGGHSVK.R (Ions score 115) |
| 243 - 254 | 1362.6762 | 1361.6690 | 1361.6728 | -3 | 0 | K.DQIFQFGGHSVK.R (No match) |
| 243 - 255 | 1518.8134 | 1517.8061 | 1517.7739 | 21 | 1 | K.DQIFQFGGHSVKR.A (No match) |
| 275 - 289 | 1610.8257 | 1609.8184 | 1609.8134 | 3 | 0 | K.AEEIGLPVHMNTTAK.N (No match) |
| 290 - 298 | 1045.5470 | 1044.5397 | 1044.5200 | 19 | 0 | K.NLIQDETGR.V (No match) |
| 318 - 333 | 1593.8105 | 1592.8032 | 1592.7803 | 14 | 0 | K.AVVMAAGGFGANIEMR.K (No match) |
| 318 - 334 | 1721.9194 | 1720.9122 | 1720.8753 | 21 | 1 | K.AVVMAAGGFGANIEMRK.R (No match) |
| 318 - 334 | 1737.8945 | 1736.8872 | 1736.8702 | 10 | 1 | K.AVVMAAGGFGANIEMRK.R Oxidation (M) (No match) |
| 335 - 343 | 1225.5722 | 1224.5650 | 1224.5523 | 10 | 1 | K.RFNPEYDER.Y (No match) |
| 336 - 343 | 1069.4804 | 1068.4731 | 1068.4512 | 20 | 0 | R.FNPEYDER.Y (No match) |
| 344 - 366 | 2220.1178 | 2219.1105 | 2219.0641 | 21 | 0 | R.YGTTNHAGATGDGIVMSAAINAK.T (No match) |
| 367 - 393 | 2785.4588 | 2784.4515 | 2784.4116 | 14 | 0 | K.TANLGEIQAYPICNPETGAIALIADAR.F (No match) |
| 367 - 393 | 2856.4861 | 2855.4788 | 2855.4487 | 11 | 0 | K.TANLGEIQAYPICNPETGAIALIADAR.F Propionamide (C) (Ions score 77) |
| 394 - 405 | 1322.6902 | 1321.6829 | 1321.7030 | -15 | 0 | R.FFGAILVNQEGK.R (No match) |
| 394 - 406 | 1478.8087 | 1477.8015 | 1477.8041 | -2 | 1 | R.FFGAILVNQEGKR.F (No match) |
| 407 - 413 | 907.4706 | 906.4633 | 906.4447 | 21 | 0 | R.FVEELDR.R (No match) |
| 407 - 414 | 1063.5592 | 1062.5520 | 1062.5458 | 6 | 1 | R.FVEELDRR.D (No match) |
| 415 - 427 | 1372.7434 | 1371.7362 | 1371.7358 | 0 | 0 | R.DVISNAILNQTGK.Y (No match) |
| 428 - 436 | 1214.6130 | 1213.6057 | 1213.6131 | -6 | 0 | K.YTYVIWNQK.I (No match) |
| 443 - 457 | 1768.8527 | 1767.8455 | 1767.7999 | 26 | 0 | K.TIDMHPGEFNDLHSR.G (Ions score 72) |
| 458 - 472 | 1623.7990 | 1622.7917 | 1622.7861 | 3 | 0 | R.GLMFEVESIEEAAAK.F (No match) |
| 473 - 495 | 2564.3383 | 2563.3310 | 2563.3282 | 1 | 1 | K.FNIPLDTLQNTIKDVNQYAATGK.D (No match) |
| 496 - 502 | 849.4341 | 848.4268 | 848.4140 | 15 | 0 | K.DLAFNNR.A (No match) |
| 513 - 517 | 722.3911 | 721.3838 | 721.4163 | -45 | 0 | K.YWILK.A (No match) |
| 535 - 546 | 1300.6861 | 1299.6788 | 1299.6670 | 9 | 0 | K.AEVLNNDGDIIK.G (No match) |
| 547 - 563 | 1702.8782 | 1701.8710 | 1701.8434 | 16 | 0 | K.GLYAAGEVTGLTHGSNR.L (No match) |
| 564 - 577 | 1495.8206 | 1494.8133 | 1494.7831 | 20 | 0 | R.LGGNAYTDIIVFGR.I (Ions score 134) |
| 578 - 595 | 2039.0408 | 2038.0336 | 2038.0007 | 16 | 1 | R.IAGQEAAKQEFEAYVEQK.L (No match) |
| 586 - 595 | 1270.5797 | 1269.5724 | 1269.5877 | -12 | 0 | K.QEFEAYVEQK.L (No match) |
| 596 - 613 | 2127.0037 | 2125.9964 | 2125.9678 | 13 | 0 | K.LISEEDLNSAVDHHHHHH.- (Ions score 88) |

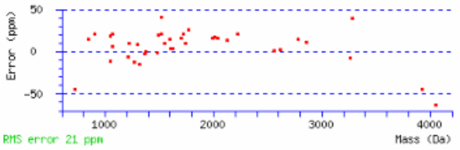


**Fig. S2.** Identification of the purified Swoo_3912 protein by MS- and MS/MS-analyses.

### {MATRIX SCIENCE} Mascot Search Results

#### Protein View

Match to: **Swoo_3913_6His** Score: **1260** Expect: **3e-122**
**Swoo_3913_6His**
Found in search of DATA.TXT

Nominal mass ($M_r$): **59034**; Calculated pI value: **5.73**
NCBI BLAST search of Swoo_3913_6His against nr
Unformatted sequence string for pasting into other applications

Variable modifications: Oxidation (M),Propionamide (C)
Cleavage by Trypsin: cuts C-term side of KR unless next residue is P
Sequence Coverage: **80%**

Matched peptides shown in **Bold Red**

```
  1 MDTISLTGKD LTIEQLLKVQ NGVASITVSQ QGKQNIQNSF DLLMDSARQG
 51 TPVYGLTVGV GKNKDTEVFG KDGKMSDEAV KLSEDFNRDM LFTHSAASGE
101 PIEQSVVRMA MTIRLNQIAT GHVGVQPDVA RLYEEFVNKN IIPVVPSDGS
151 VGLSDILLAS HIGAAMMGEH EVFYKGERMP AGKALKAAGV KPLVPFGKDG
201 LSILSNNALS TAQLVHAMGQ AEQLIEFSPK LIAAGLESIN GNISPILPHT
251 LDARPMPHVQ EVGNDILTNL EGGYLFQRDE KRPLQDALSF RGAHWPVANA
301 ISQYNSLEEL VHIQINSSDD NPTVYLNAKH SRFNKYPQVE QYFTDGKVSG
351 AINSSANFDT IQLATTTEAF SIAMAQLGKY SSNRQIKMID PYFTGLPRAL
401 VNPDDTNGQS FYTLQNGFTA LFVDIAHASN PVSFYGQANQ GGIEDNFSNF
451 HQASKNLTTV VNGVSYIYAF ELMTYGQALD LQKKVHNRDL SDANKTLLKD
501 LRKTVDFYAD DTRTFTIDIE ASQKFLMKYL EHHHHHH
```

Show predicted peptides also

Sort Peptides By ◉ Residue Number ○ Increasing Mass ○ Decreasing Mass

| Start - End | Observed | Mr(expt) | Mr(calc) | ppm | Miss | Sequence |
|---|---|---|---|---|---|---|
| 10 - 18 | 1072.6296 | 1071.6223 | 1071.6176 | 4 | 0 | K.DLTIEQLLK.V (No match) |
| 19 - 33 | 1515.8440 | 1514.8367 | 1514.8053 | 21 | 0 | K.VQNGVASITVSQQGK.Q (Ions score 98) |
| 34 - 48 | 1751.8461 | 1750.8388 | 1750.8308 | 5 | 0 | K.QNIQNSFDLLMDSAR.Q (Ions score 113) |
| 49 - 62 | 1375.7529 | 1374.7456 | 1374.7507 | -4 | 0 | R.QGTPVYGLTVGVGK.N (No match) |
| 63 - 71 | 1037.5346 | 1036.5273 | 1036.5189 | 8 | 1 | K.NKDTEVFGK.D (No match) |
| 75 - 81 | 779.3603 | 778.3530 | 778.3531 | -0 | 0 | K.MSDEAVK.L (No match) |
| 82 - 88 | 880.4169 | 879.4097 | 879.4086 | 1 | 0 | K.LSEDFNR.D (No match) |
| 82 - 108 | 3035.5218 | 3034.5145 | 3034.4454 | 23 | 1 | K.LSEDFNRDMLFTHSAASGEPIEQSVVR.M (No match) |
| 89 - 108 | 2174.1068 | 2173.0995 | 2173.0474 | 24 | 0 | R.DMLFTHSAASGEPIEQSVVR.M (Ions score 134) |
| 109 - 114 | 722.3480 | 721.3407 | 721.3615 | -29 | 0 | R.MAMTIR.L (No match) |
| 115 - 131 | 1775.0115 | 1774.0042 | 1773.9486 | 31 | 0 | R.LNQIATGHVGVQPDVAR.L (Ions score 80) |
| 132 - 139 | 1041.5232 | 1040.5159 | 1040.5178 | -2 | 0 | R.LYEEFVNK.N (No match) |
| 140 - 175 | 3766.9249 | 3765.9176 | 3765.9110 | 2 | 0 | K.NIIPVVPSDGSVGLSDILLASHIGAAMMGEHEVFYK.G (Ions score 133) |
| 140 - 178 | 4108.9910 | 4107.9837 | 4108.0762 | -23 | 1 | K.NIIPVVPSDGSVGLSDILLASHIGAAMMGEHEVFYKGER.M (No match) |
| 187 - 198 | 1183.6968 | 1182.6895 | 1182.7125 | -19 | 0 | K.AAGVKPLVPFGK.D (No match) |
| 199 - 230 | 3382.7736 | 3381.7664 | 3381.7238 | 13 | 0 | K.DGLSILSNNALSTAQLVHAMGQAEQLIEFSPK.L (Ions score 138) |
| 279 - 291 | 1574.8591 | 1573.8518 | 1573.8212 | 19 | 1 | R.DEKRPLQDALSFR.G (No match) |
| 282 - 291 | 1202.6504 | 1201.6432 | 1201.6567 | -11 | 0 | K.RPLQDALSFR.G (No match) |
| 292 - 329 | 4207.9533 | 4206.9460 | 4207.0610 | -27 | 0 | R.GAHWPVANAISQYNSLEELVHIQINSSDDNPTVYLNAK.H (Ions score 161) |
| 333 - 347 | 1863.8835 | 1862.8762 | 1862.8839 | -4 | 1 | R.FNKYPQVEQYFTDGK.V (No match) |
| 348 - 379 | 3257.6590 | 3256.6518 | 3256.6285 | 7 | 0 | K.VSGAINSSANFDTIQLATTTEAFSIAMAQLGK.Y (No match) |
| 380 - 384 | 626.2395 | 625.2322 | 625.2820 | -80 | 0 | K.YSSNR.Q (No match) |
| 380 - 387 | 995.5147 | 994.5075 | 994.5196 | -12 | 1 | K.YSSNRQIK.M (No match) |
| 388 - 398 | 1309.6604 | 1308.6531 | 1308.6536 | -0 | 0 | K.MIDPYFTGLPR.A (No match) |
| 399 - 455 | 6162.8624 | 6161.8551 | 6162.8685 | -164 | 0 | R.ALVNPDDTNGQSFYTLQNGFTALFVDIAHASNPVSFYGQANQGGIEDNFSNFHQASK.N (No match) |
| 456 - 483 | 3151.6361 | 3150.6289 | 3150.5947 | 11 | 0 | K.NLTTVVNGVSYIYAFELMTYGQALDLQK.K (Ions score 199) |
| 514 - 524 | 1252.6471 | 1251.6398 | 1251.6347 | 4 | 0 | R.TFTIDIEASQK.F (No match) |
| 529 - 537 | 1246.6030 | 1245.5958 | 1245.5540 | 34 | 0 | K.YLEHHHHHH.- (No match) |

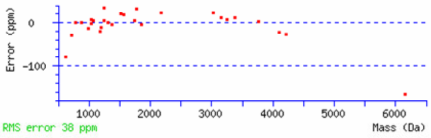


**Fig. S3.** Identification of the purified Swoo_3913 protein by MS- and MS/MS-analyses.

```
---
So_UrdA  : --MHYKKSIIGIAVTATAIIAGCQVTHQIVKSQGTAQGKHGEVQVETTFKDGHIVAIDVLKQKENKVLAGAVFKDVKQAIIDNNSIEVDGIAGATVTSKALKEAVGKSIEAAGVTLVATASAK-KSEALTPAEYTYD : 134
Swoo_3912: MNFKFKKSVLGMAVAVPVVLVGCAVTNEMVKSQGTAKGRHGDITVETTFENGKITAIDIVKQKENRVLSAAVYKDVKQAIIDNNSINVDGITGATATSDGLKKAVAKSAELAGITLVATAAINGKKVNAEPSEYTYD : 137

So_UrdA  : VVIIGSGGAGFSAGLEAIAAGRSAVIIEKMPIIGGNSLISGAEMNVAGSWVQKNMGITDSKELFISDTLKGGDFKGDPEMVKTMVDNAVGAAEWLRDYVKVEFYPDQLFQFGGHSVKRALIPKGHTGAEVISKFSIK : 271
Swoo_3912: VVVIGAGGAGFSAGVEAVEAGVSAVIIEKQPIIGGNSLISGGEMNVAGSWVQQSMGITDSKALFIEDTLKGGDYKGDPEMVRVMADNAVAAAEWLRDDIKVDFYKDQIFQFGGHSVKRAVIPKGHTGAEVLSKFAAK : 274

So_UrdA  : ADEVGLPIHTNTKAEKLIQDQTGRIVGVEAAHNGKTITYHAKRGVVIATGGFSSNMEMRKKYNPELDERYGSTGHAGGTGDGIVMAEKIHAAAKNMGYIQSYPICSPTSGAIALIADSRFFGAVLINQKGERFVEEL : 408
Swoo_3912: AEEIGLPVHMNTTAKNLIQDETGRVVGVKAMKNGKVITYHAKKAVVMAAGGFGANIEMRKRFNPEYDERYGTTNHAGATGDGIVMSAAINAKTANLGEIQAYPICNPETGAIALIADARFFGAILVNQEGKRFVEEL : 411

So_UrdA  : ERRDVISHAILAQPGRYTYVLWNQDIENVAHTVEMHQGELKEFTKDGLMYKVDTLEEAAKVFNIPEDKLLSTIKDVNHYAATGKDEAFNHRSGLVDLSKGPYWILKATPSVHHTMGGLVVDTRTRVLDEQGKVIPGL : 545
Swoo_3912: DRRDVISNAILNQTGKYTYVIWNQKIDDLAKTIDMHPGEFNDLHSRGLMFEVESIEEAAAKFNIPLDTLQNTIKDVNQYAATGKDLAFNNRAGLVDMSEGKYWILKATPSVHHTMGGIATTTKAEVLNNDGDIIKGL : 548

So_UrdA  : FAAGEVTGLTHGTNRLGGNAYTDIIVYGRIAGQEAAK- : 582
Swoo_3912: YAAGEVTGLTHGSNRLGGNAYTDIIVFGRIAGQEAAKQ : 586
```

**Fig. S4.** Sequence alignment of *S. oneidensis* urocanate reductase (*So*_UrdA, UniProt Id: Q8CVD0) and *S. woodyi* Swoo_3912 protein (UniProt Id: B1KFJ5). The residues involved in urocanate carboxylic group and N-atom binding are marked by red and blue color, respectively. Lipidation site is marked by yellow. Flavinylation motif is marked by magenta with FMN attachment site colored in green.

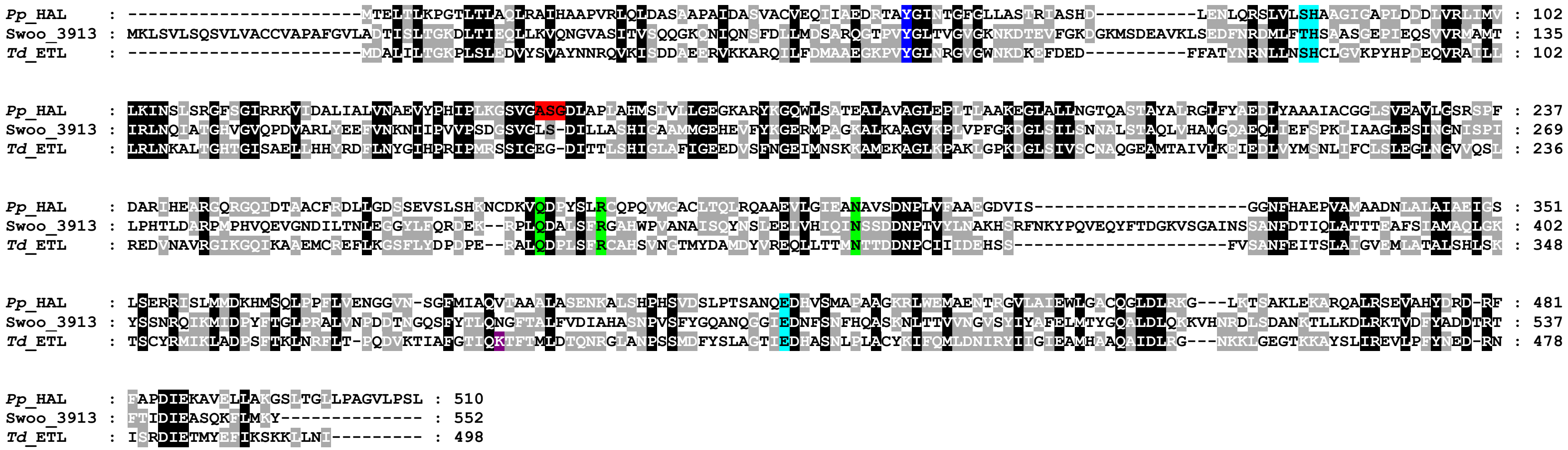

**Fig. S5.** Multiple sequence alignment of histidine ammonia-lyase from *P. putida* (*Pp*_HAL, GenBank Id: BAN56952), ergothionase from *T. denticola* (*Td*_ETL, GenBank Id: EMB27117), and the Swoo_3913 protein from *S. woodyi* (GenBank Id: ACA88171). Residues involved in the binding of the substrate carboxylic group and imidazole N-atoms are marked by green and light-blue color, respectively. Tyr residues essential for proton withdrawal from substrate β-position are marked by blue. Ala-Ser-Gly sequence, forming the MIO-group of ammonia-lyases, is colored in red. Lys384 essential for the binding of the thiolate group of ergothioneine in *Td*_ETL is marked by magenta.

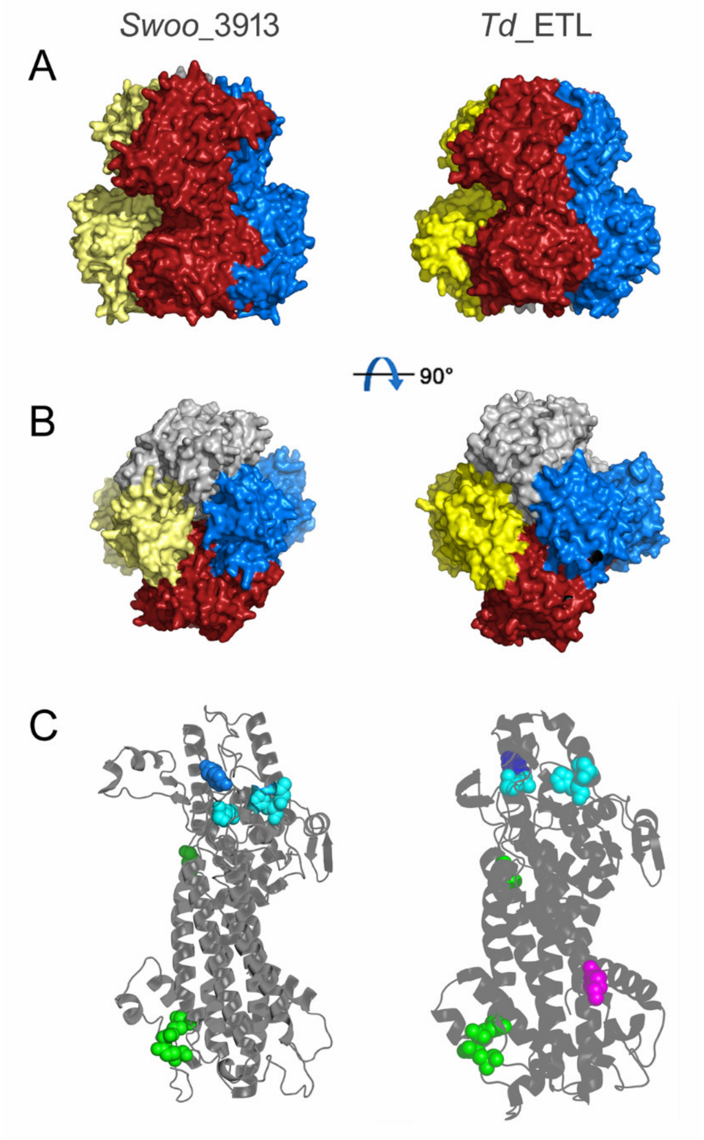


**Fig. S6.** Comparison of the 3D structures of the *S. woodyi* Swoo_3913 protein (predicted by AlphaFold 3) and *T. denticola* ergothionase *Td*_ETL (crystal structure; PDB Id: 6s7j). **A** and **B**, Two views of protein homotetramers with differently colored single-domain subunits. **C**, An enlarged view of separate "grey" subunit from panel A shown in the same orientation in a cartoon representation. Several residues shown as spheres are those highlighted in the same colors in Fig, S5. These residues belong to *Td*_ETL active sites, each of which is formed by residues from three subunits. The figure was created using Pymol (PyMOL Molecular Graphics System, version 1.5.0.4; Schrödinger, LLC).

### Supplementary methods

##### Histidine betaine (HB) synthesis

HB was synthesized from L-histidine by methylation with iodomethane [Valeev et al., 2007]. L-Histidine (2.24 g) was dissolved in 30 mL of a water/methanol (2:1, v/v) mixture. The pH value was adjusted to 11 with 3 M NaOH, 7.12 g of iodomethane were added with stirring, and the resulting mixture was stirred at room temperature while maintaining pH 11 with 3 M NaOH portions until the pH stabilized (~6 h). Then, the mixture was neutralized with hydrochloric acid and loaded on a column of Dowex 50-X4 resin (2.6 × 22 cm) in the $H^+$ form. The column was washed with water, and HB was eluted with 2 M HCl solution. Aliquots of the eluate were tested with the Pauly's diazo test [Bogachev, 2013]. The diazo-positive eluate fractions were combined and evaporated to dryness under reduced pressure over KOH flakes. The residue was extracted with hot methanol, and HB was crystallized by the addition of diethyl ether. HB was further recrystallized from methanol twice by the addition of diethyl ether. The final product contained 95% HB, as determined by mass spectrometry.
